# Baseline Predictors of Craving Reduction Following Non-Invasive Neuromodulation: A Computational Modeling Approach

**DOI:** 10.64898/2026.11.18.767490

**Authors:** Jungyoon Kang, Jung-Seok Choi, Young-Chul Jung, Minjae Kim, Minkyung Park, Manjae Kwon, Yunesang Lee, Dongil Chung, Woo-Young Ahn

## Abstract

**Background:** Alcohol craving is a clinically relevant symptom with an established neurobiological basis and has emerged as a treatment target. Non-invasive neuromodulation has shown potential for reducing craving, but substantial inter-individual variability in response is a pervasive challenge. This study examined whether baseline computational parameters predicted treatment response following neuromodulation.

**Methods:** This secondary analysis of an open-label, randomized trial included 58 individuals with subclinical alcohol use who received transcutaneous auricular vagus nerve stimulation (taVNS), transcranial photobiomodulation (tPBM), or combined taVNS + tPBM for five weeks. Baseline computational measures from Delay Discounting, Choice under Risk and Ambiguity, and Loss Aversion Tasks were examined as predictors of treatment response. Elastic net regularization was used to identify markers associated with Reliable Change Index (RCI)-defined responder status, alongside demographic variables, intervention condition, and baseline craving.

**Results:** Of the 58 participants, 24 were classified as treatment responders. The elastic net model showed a mean test AUC of 0.66 across 1,000 iterations, indicating modest predictive performance. Baseline craving, intervention condition, age, sex, and computational modeling parameters contributed to responder status: Higher loss aversion and lower risk sensitivity were associated with a greater likelihood of responder status.

**Conclusions:** Baseline computational measures, particularly loss aversion and risk sensitivity, may serve as candidate behavioral markers associated with craving reduction following non-invasive neuromodulation. Given the open-label design, these findings represent an initial proof-of-concept. Future sham-controlled studies are needed to determine whether these parameters are associated with stimulation-specific responses or more general susceptibility to behavioral change.

**Clinical trial registration:** The trial was registered at ClinicalTrials.gov (NCT07071493).

## 1. Introduction

Alcohol craving is a clinically relevant feature of alcohol use that is closely associated with drinking behavior across the continuum of alcohol involvement (Yoon et al., 2021; Grüsser et al., 2006). It arises from interactions among neural systems involved in reward processing, affect regulation, and cognitive control (Koob and Volkow, 2010, 2016; Kober and Mell, 2015). Because modulating these systems has been shown to reduce craving and drinking behavior (D. J. Kim et al., 2025), non-invasive neuromodulation has emerged as a promising therapeutic approach.

Specifically, non-invasive neuromodulation can influence craving-relevant neural processes through distinct routes. Peripheral approaches, such as transcutaneous auricular or cervical vagus nerve stimulation (taVNS, tcVNS), engage vagal afferents that project to brainstem-cortical networks involved in affective and stress regulation (Thompson-Lake et al., 2026; Frangos & Komisaruk, 2017); both forms of VNS have been reported to improve alcohol-related depressive symptoms (Klaming et al., 2025; Wang et al., 2021), although their effects on craving itself remain unclear (Kwon et al., 2026; Wang et al., 2021). Transcranial modalities— including electrical stimulation (tES), magnetic stimulation (TMS), and photobiomodulation (tPBM)—can be applied to cortical targets, particularly prefrontal regions implicated in cognitive control (O’Connor et al., 2025; Soleimani et al., 2024); recent meta-analyses support the efficacy of tES and TMS for alcohol craving, with effects varying by stimulation modality and protocol (D. J. Kim et al., 2025; Mehta et al., 2024), and tPBM was recently shown to produce significant alcohol craving reduction (Kwon et al., 2026).

Although these findings support the potential of non-invasive neuromodulation for reducing alcohol craving, group-level efficacy does not imply uniform treatment response. Indeed, substantial inter-individual variability in response is a pervasive challenge across non-invasive neuromodulation approaches (Soleimani et al., 2023). This variability may partly reflect the fact that similar clinical presentations, such as alcohol craving, can arise from distinct cognitive and neurobiological processes (Kwako et al., 2016; Redish et al., 2008), which may in turn produce heterogeneous responses to a given treatment. Accordingly, efforts to account for this variability have examined pretreatment predictors across multiple domains. Clinical and conventional cognitive measures are readily accessible and have shown predictive value (Chen et al., 2020; Ren et al., 2024; Toffanin et al., 2022), but may provide limited process-level specificity. Neurophysiological and neuroimaging measures can provide more direct insight into stimulation-relevant neural function (Ghazi et al., 2025; Klooster et al., 2024; Zhao et al., 2024), but their cost, technical complexity, and analytic demands may limit routine clinical use (Constantinides et al., 2026; Hernandez-Pavon et al., 2023). Thus, existing approaches involve a trade-off between process specificity and practical scalability, highlighting the need for pretreatment markers that can offer both.

Model-based computational modeling parameters may help bridge this gap by quantifying latent decision processes from behavioral assessments. By fitting formal mathematical models to choices in cognitive tasks, computational modeling can separate distinct decision processes that may otherwise produce similar observable behavior and estimate these processes at the individual level (Farrell and Lewandowsky, 2018; Montague et al., 2012). Importantly, many of these decision processes are supported by neural systems involved in valuation, affective processing, and cognitive control (Hiser & Koenigs, 2018; Huys et al., 2016; Rangel et al., 2008), which also contribute to craving-related behavior (Koob and Volkow, 2010, 2016). This provides a mechanistic rationale for examining baseline computational measures as potential markers of neuromodulation response, given that the effects of stimulation may depend partly on the pre-existing functional state of the targeted systems (Bradley et al., 2022; Krause & Cohen Kadosh, 2014). However, such computational measures do not directly index these systems or neural target engagement. Rather, they quantify baseline individual differences in latent decision processes that may shape an individual’s capacity for behavioral change following neuromodulation.

Emerging evidence suggests that computational modeling parameters can have prognostic value in substance-use treatment more broadly. Parameters reflecting risk-taking have been linked to subsequent relapse among individuals receiving treatment for alcohol use disorder (AUD) (Yuan et al., 2024; Sasaki et al., 2023), while delay discounting has yielded mixed evidence—predictive of smoking-cessation outcomes in a recent study (J.-H. Lee et al., 2026) but inconsistent in a broader systematic review (Exum et al., 2023). Other parameters, including loss aversion (Todesco et al., 2022) and ambiguity sensitivity (J.-H. Lee et al., 2026; Konova et al., 2020), have also been implicated in substance-use phenotypes and treatment-related dynamics, although their value as baseline predictors remains less established, warranting investigation in novel interventions such as neuromodulation.

Prior studies have used computational models to quantify neuromodulation-induced changes in specific decision processes (e.g., Nejati et al., 2025; Liu et al., 2024; Filmer et al., 2020), but to our knowledge, none have examined pretreatment computational measures as predictive markers of subsequent craving reduction. In this secondary analysis of a randomized neuromodulation trial in individuals with subclinical alcohol use, we used an experimental optimization technique to efficiently and reliably estimate individual-level decision-making parameters and examine four baseline parameters —delay discounting rate, loss aversion, risk sensitivity, and ambiguity sensitivity—as predictors of treatment response using elastic net logistic regression. Treatment response was defined as a reliable reduction in alcohol craving, based on the Reliable Change Index (RCI), following five weeks of taVNS, tPBM, or their combination. Given prior evidence linking greater risk-taking tendencies to poorer alcohol-related outcomes (Yuan et al., 2024; Sasaki et al., 2023), we hypothesized that lower risk sensitivity (i.e., lower risk-taking tendency) would be associated with a greater likelihood of responder classification. The remaining parameters were examined exploratorily, without directional hypotheses. As an exploratory aim, we also examined pre–post changes in these computational measures within each intervention group.

## 2. Methods

### 2.1 Participants and Study Design

This study reports a secondary analysis of data collected from an open-label, randomized, three-arm parallel-group trial examining the effects of non-invasive neuromodulation on alcohol craving (Kwon et al., 2026). Participants provided written informed consent prior to participation. The study was approved by the Institutional Review Board of the participating institution (IRB No. 4-2024-1389) and was conducted in accordance with the Declaration of Helsinki. The trial was registered at ClinicalTrials.gov (NCT07071493).

Inclusion criteria were as follows: adults aged 19 to 40 years; current alcohol drinkers; capacity to provide valid informed consent; a score of 4 or higher on the Korean version of the Alcohol Use Disorders Identification Test (AUDIT) (Lee, 2000); absence of DSM-5 criteria for AUD; and ability to complete the five-week stimulation program. Exclusion criteria were as follows: inability to read or comprehend the consent form; lack of capacity for independent decision-making; vulnerable status, including pregnancy or affiliation with the research institution as an employee, student, investigator, or study sponsor; current substance use disorder involving substances other than alcohol or nicotine; and metal allergy that could interfere with stimulation procedures.

Participants were randomly assigned to one of three intervention groups: taVNS, tPBM, or combined taVNS + tPBM. Clinical and behavioral assessments were conducted at baseline (T0) and following the five-week intervention period (T1). The overall study design is illustrated in **Figure 1**. Of the 60 participants initially enrolled, one withdrew during the intervention due to a headache, and one did not complete the baseline survey, resulting in a final analytic sample of 58 participants (Table 1).

**Figure 1.**
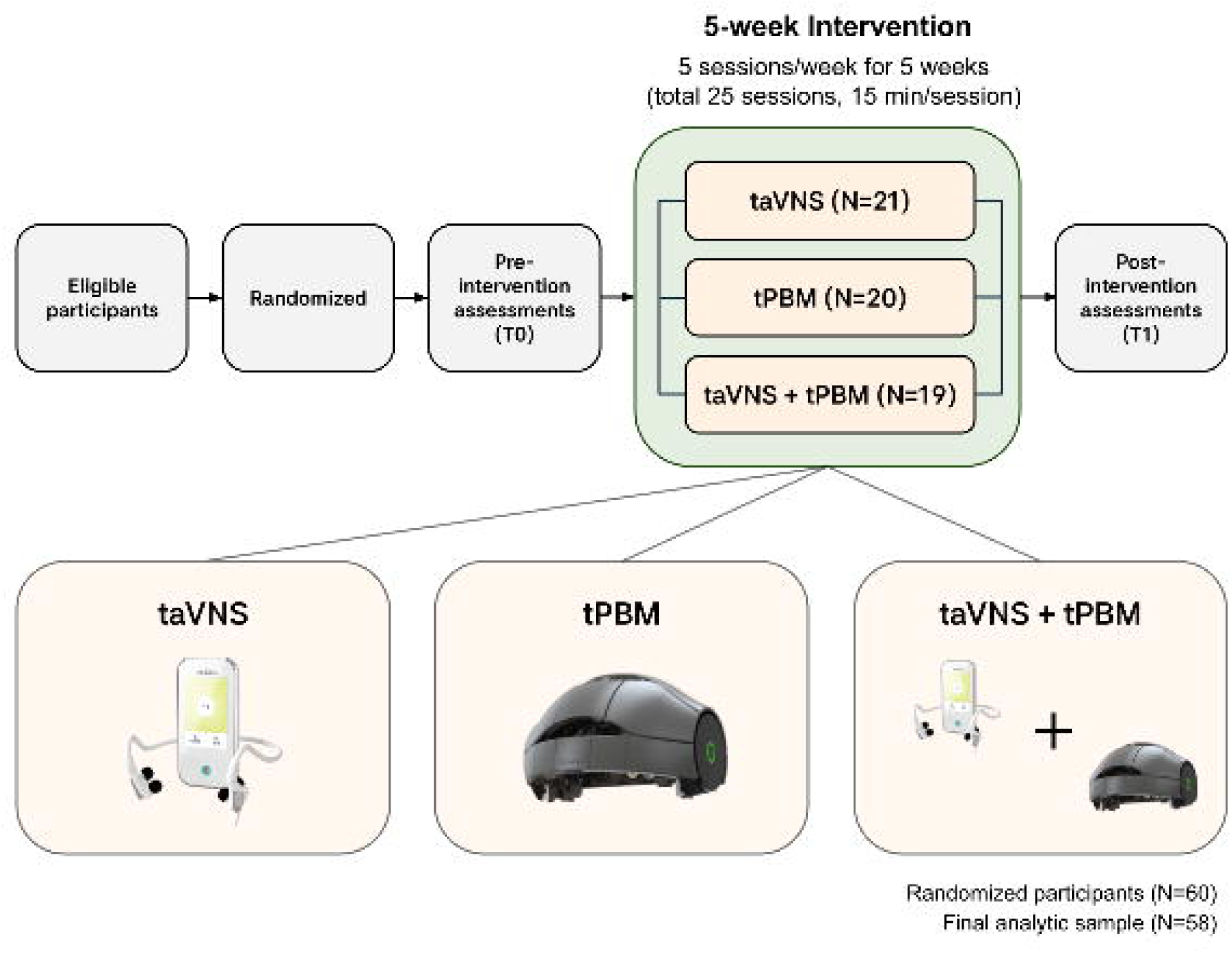
Overview of the study design. Participants were randomized to one of three intervention arms (taVNS, tPBM, or combined taVNS + tPBM) and assessed at baseline (T0) and post-intervention (T1). Of 60 randomized participants, 58 were retained for the final analysis.

**Table 1.** Baseline demographic, clinical, and computational characteristics of participants in the three intervention groups. Group differences were assessed using one-way ANOVA for continuous variables and the chi-square test for categorical variables. taVNS, transcutaneous auricular vagus nerve stimulation; tPBM, transcranial photobiomodulation; PACS, Penn Alcohol Craving Scale; AUDIT, Alcohol Use Disorders Identification Test; DDT, Delay Discounting Task; LA, Loss Aversion Task; CRA, Choice under Risk and Ambiguity Task. **Note**. CRA risk sensitivity (α) is reported on its original scale in Table 1; the rescaled parameter (α − 1) was used in the elastic net analyses to facilitate interpretation.

| **Characteristics** | **taVNS<br>(N=20)** |  | **tPBM<br>(N=19)** |  | **taVNS + tPBM<br>(N=19)** |  | **p-value** |
|---|---|---|---|---|---|---|---|
|  | Mean (N) | SD (%) | Mean (N) | SD (%) | Mean (N) | SD (%) |  |
| **Demographic** |  |  |  |  |  |  |  |
| **Age** | 26.45 | 4.51 | 28.53 | 4.89 | 27.63 | 4.79 | 0.394 |
| **Sex (Male, %)** | 9 (45.0%) |  | 8 (42.1%) |  | 8 (42.1%) |  | 0.978 |
| **Marriage (Unmarried, %)** | 18 (90.0%) |  | 17 (89.5%) |  | 16 (84.2%) |  | 0.830 |
| **Education (Years)** | 15.65 | 1.42 | 15.84 | 1.95 | 16.21 | 1.99 | 0.620 |
| **PACS** | 7.95 | 5.34 | 7.32 | 4.71 | 6.42 | 2.67 | 0.558 |
| **AUDIT** | 10.55 | 6.16 | 10.58 | 5.01 | 10.37 | 5.20 | 0.992 |
| **Computational parameters** |  |  |  |  |  |  |  |
| **DDT discounting rate (log (k))** | -3.55 | 1.57 | -4.52 | 1.40 | -4.05 | 1.07 | 0.098 |
| **LA loss aversion (λ)** | 1.86 | 0.92 | 2.12 | 0.70 | 1.81 | 0.83 | 0.450 |
| **CRA risk sensitivity (α)** | 0.55 | 0.30 | 0.55 | 0.22 | 0.58 | 0.27 | 0.919 |
| **CRA ambiguity sensitivity (β)** | 0.73 | 0.69 | 0.60 | 0.47 | 0.63 | 0.68 | 0.799 |

### 2.2 Neuromodulation Conditions

Participants self-administered one of three interventions (taVNS, tPBM, or combined taVNS + tPBM) remotely over five weeks, with five 15-minute sessions per week. Detailed stimulation parameters and device specifications are provided in the Supplementary Methods.

### 2.3 Behavioral Tasks and Computational Modeling

Baseline computational modeling parameters were estimated from three decision-making tasks: the Delay Discounting Task (DDT) (Green & Myerson, 2004), the Choice under Risk and Ambiguity Task (CRA) (Levy et al., 2010), and the Loss Aversion (LA) Task (Tom et al., 2007). Tasks were administered at both T0 and T1, but only baseline (T0) parameters were used as predictors in the present analyses, as our goal was to identify pre-intervention computational markers predictive of subsequent treatment response.

The three tasks involve a binary choice on every trial, as illustrated in **Figure 2**. All behavioral tasks were implemented using ADO, a Bayesian approach in which posterior distributions over model parameters were updated after each choice using Bayesian grid inference with uniform priors (Yang et al., 2021; Myung et al., 2013). This adaptive procedure selected informative trials based on participants’ previous choices, enabling efficient and reliable estimation of individual-level computational parameters (Ahn et al., 2020; Kwon et al., 2023; J.-H. Lee et al., 2026). ADO-based tasks were performed using a Python library called ADOpy (Yang et al., 2021).

**Figure 2.**
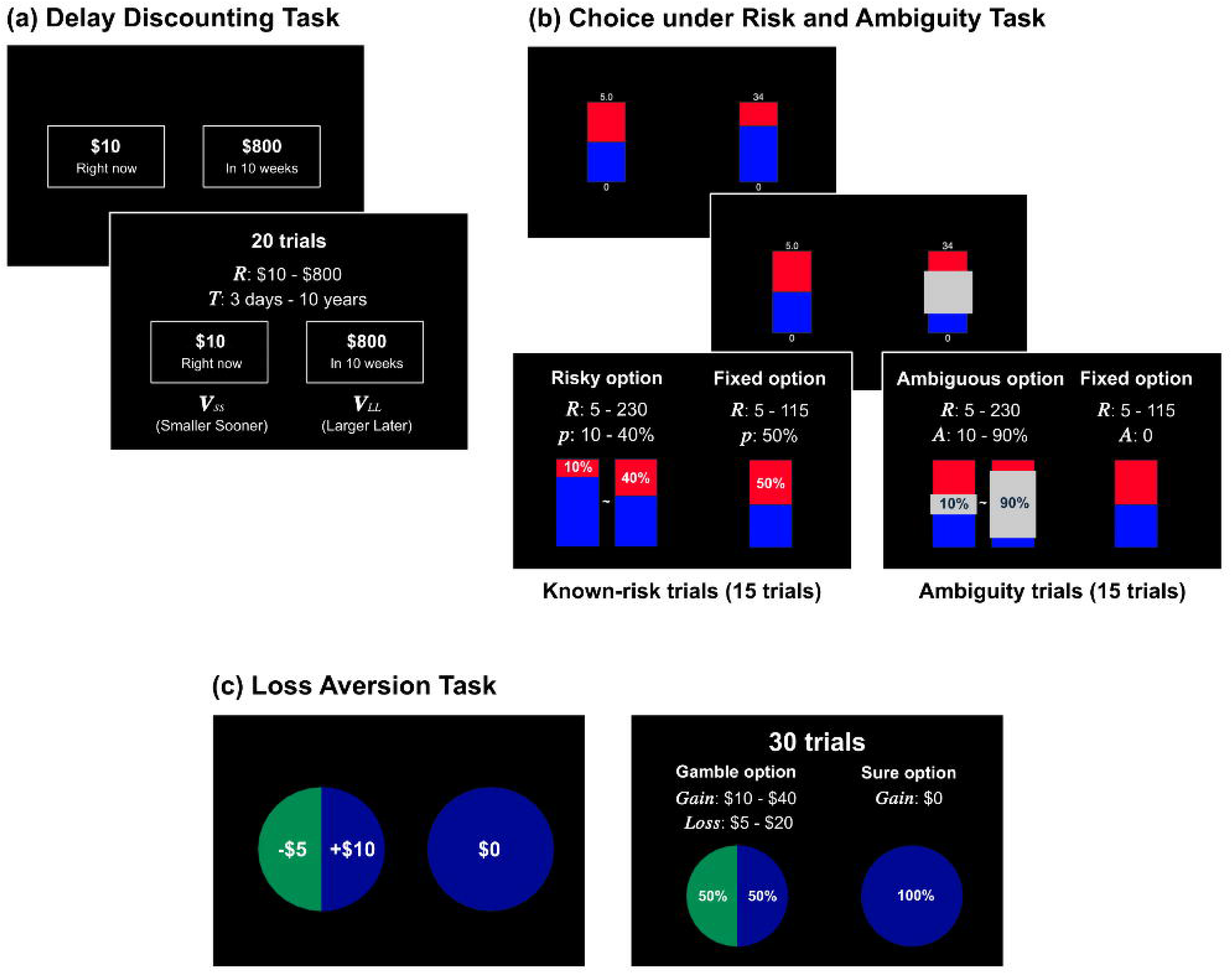
Overview of the three decision-making tasks and their computational parameters. (a) The Delay Discounting Task (DDT) involved choices between smaller immediate and larger delayed rewards and yielded discounting rate (k). (b) The Choice under Risk and Ambiguity Task (CRA) involved choices between fixed and variable options under known-risk and ambiguous probability conditions and yielded risk sensitivity (α) and ambiguity sensitivity (β). (c) The Loss Aversion Task (LA) involved accept/reject decisions for mixed gambles and yielded loss aversion (λ). Dollar signs are shown to aid interpretation, although participants completed the task using Korean won amounts on a comparable scale.

#### 2.3.1 Computational models and parameters in the DDT (Green & Myerson, 2004)

In the DDT, participants chose between a smaller immediate reward (Smaller Sooner, SS) and a larger delayed reward (Larger Later, LL) (Figure 2a). The task consisted of 20 trials, and choices were used to estimate the discounting rate, which quantifies the degree to which delayed rewards were devalued. The discounting rate was estimated using a hyperbolic model,

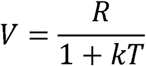

where *V* denotes the subjective value of each option (computed separately for SS and LL), *R* the reward amount, *κ* the discounting rate, and *T* denotes the delay. Higher k values indicate steeper discounting of delayed rewards, reflecting a stronger preference for immediate over delayed rewards.

Choice probabilities were modeled using a softmax function:

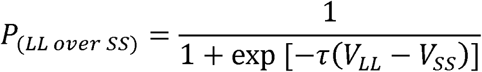

where *τ* denotes the inverse temperature parameter. Given the skewed distribution of the discounting rate k, a log-transformation was applied for subsequent analyses.

#### 2.3.2 Computational models and parameters in the CRA (Levy et al., 2010)

The CRA consisted of 30 trials, including equal numbers of risk and ambiguity trials. In each trial, participants chose between a fixed option, offering a smaller reward with a known 50% probability of receipt, and a variable option offering a larger reward with either a known (risk trials) or partially hidden (ambiguity trials) probability of receipt; full stimulus ranges are shown in Figure 2b.

Two computational modeling parameters, risk sensitivity and ambiguity sensitivity, were estimated from this task using a modified utility model (Levy et al., 2010). In this model, the subjective utility of each option was determined by its reward amount, reward probability, and ambiguity level, and was defined as follows:

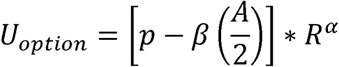

where *ρ* denotes the reward probability, *β* ambiguity sensitivity, *A* the ambiguity level (set to 0 in risk trials), *R* the reward amount, and *α* risk sensitivity, computed separately for the fixed and variable options. Values of *α* greater than 1 indicate risk-seeking, whereas values less than 1 indicate risk aversion. A value of *α* = 1 indicates risk neutrality. For *β*, positive values indicate ambiguity aversion, while negative values indicate ambiguity tolerance (tolerance for unknown probabilities). A value of *β* = 0 indicates ambiguity neutrality. To facilitate interpretation in subsequent analyses, *α* was rescaled as *α* – 1, such that positive values indicate risk-seeking and negative values indicate risk aversion. Choice probabilities were modeled using a softmax function, as described for the DDT.

#### 2.3.3 Computational models and parameters in the LA (Tom et al., 2007)

In the LA, participants decided whether to accept or reject a gamble offering a 50% chance of gain and a 50% chance of loss, relative to a sure option involving neither (30 trials; full stimulus ranges in Figure 2c).

The loss aversion parameter was estimated from participants’ choices using a prospect theory model (Sokol-Hessner et al., 2009). In this model, the subjective utility of each option was computed separately for the gain outcome, the loss outcome, the sure option, and the gamble option:

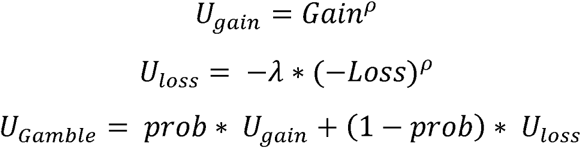

where *U_gain_* and *U_toss_* denote the subjective utilities of gain and loss outcomes, respectively, *U_sure_* the utility of the sure option, and *U_Gamble_* the expected utility of the gamble option (relative to a sure option fixed at zero utility). The risk sensitivity parameter ρ was fixed at 1 in this task. Higher *λ* values indicate greater sensitivity to loss relative to gain. Choice probabilities were modeled using a softmax function.

### 2.4 Outcome Measure

Alcohol craving was assessed at T0 and T1 as the primary outcome using the Penn Alcohol Craving Scale (PACS) (Kim et al., 2008). Treatment response was defined using the Reliable Change Index (RCI; Jacobson & Truax, 1991), which evaluates whether an individual’s pre–post change exceeds that expected from measurement unreliability. An RCI > 1.96 indicated a reliable reduction in craving, corresponding to a >3.64-point reduction from T0 to T1 in the present sample; participants meeting this criterion were classified as responders. Details of the RCI calculation are provided in the Supplementary Methods.

### 2.5 Statistical Analysis

To identify baseline predictors of RCI-defined treatment response, we fitted a penalized logistic regression model with elastic net regularization. Given the modest sample size and the potential correlations among predictors, elastic net was chosen because it can shrink regression coefficients while also selecting the most informative variables, thereby reducing overfitting and improving model stability (Zou & Hastie, 2005). RCI-defined responder status was the binary outcome (responder = 1, non-responder = 0). The four baseline computational modeling parameters were included as predictors, while baseline PACS score, age, and sex were included as covariates. Intervention condition was included as a covariate using two binary indicators (taVNS, tPBM), coded as (1, 0) for taVNS group, (0, 1) for tPBM group, and (1, 1) for combined taVNS + tPBM group, to account for previously reported differences in treatment efficacy across intervention arms (Kwon et al., 2026). All continuous predictors were standardized prior to model fitting.

Using the easyml package in R (Hendricks & Ahn, 2017), hyperparameters were explored by fixing each α value from 0.1 to 0.9 and determining the optimal λ via 10-fold cross-validation across 100 iterations, each with a random 80/20 train-test split. As model performance was comparable across α values, α = 0.1 was chosen to retain a broader set of candidate predictors. The final model was run for 1,000 iterations with the selected α to ensure stable estimation of performance, evaluated as the area under the receiver operating characteristic curve (AUC).

In addition to the classification model, pre–post changes in computational measures were examined within each intervention group as an exploratory analysis.

## 3. Results

Of the 60 participants initially enrolled in the source trial, 58 were included in this secondary analysis: taVNS group (n = 20), tPBM group (n = 19), and combined taVNS + tPBM group (n = 19). Baseline demographic, clinical, and computational decision-making parameters did not differ significantly among groups (Table 1). As reported by Kwon et al. (2026), craving significantly decreased in the tPBM and combined taVNS + tPBM groups, but not in the taVNS group.

Twenty-four participants (41.4%) were classified as responders and 34 (58.6%) as non-responders. Across intervention groups, the responders were distributed as follows: taVNS group (n = 5), tPBM group (n = 9), and combined taVNS + tPBM group (n = 10).

The elastic net model showed a mean test AUC of 0.66 (SD = 0.13) across 1,000 iterations (mean training AUC = 0.83, SD = 0.04). Coefficients were summarized as the mean and 95% CI across iterations, and predictors whose CIs excluded zero were interpreted as stable contributors (Figure 3).

**Figure 3.**
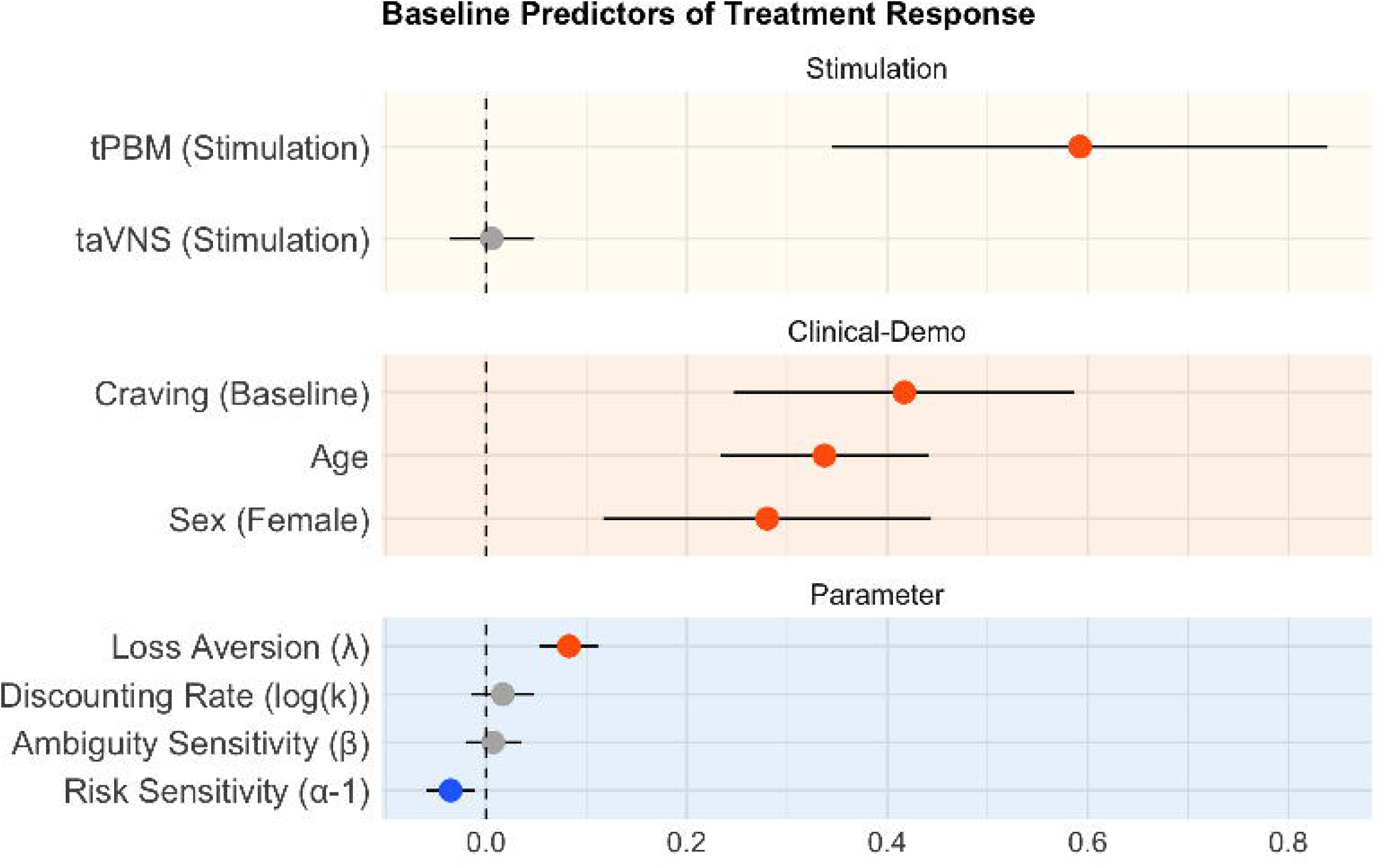
Baseline predictors of RCI-defined treatment response from elastic net logistic regression. Points represent standardized mean coefficients across 1,000 iterations with 95% confidence intervals (CI) (error bars); red and blue indicate positive and negative coefficients, respectively. Mean test AUC was 0.66 (SD = 0.13).

Among intervention conditions, tPBM showed a relatively large positive contribution to responder classification (standardized mean β coefficient = 0.59, 95% CI [0.35, 0.84]), whereas taVNS did not emerge as a stable predictor (β = 0.01, 95% CI [-0.04, 0.05]). Baseline craving score (β = 0.42, 95% CI [0.25, 0.59]), age (β = 0.34, 95% CI [0.23, 0.44]), and sex (β = 0.28, 95% CI [0.12, 0.44]) were also positively associated with treatment response, indicating that participants with higher baseline craving, older age, and female sex were more likely to respond. Among the four computational modeling parameters, loss aversion (λ; β = 0.08, 95% CI [0.05, 0.11]) and risk sensitivity (α − 1; β = −0.04, 95% CI [−0.06, −0.01]) contributed as candidate predictors. Specifically, higher loss aversion was associated with a greater likelihood of treatment response, whereas higher risk sensitivity (i.e., more positive α – 1 values, indicating a risk-seeking tendency) was negatively associated with response. The remaining computational measures, including discounting rate (log (k)) and ambiguity sensitivity (β), did not emerge as stable predictors.

As an exploratory analysis, loss aversion significantly decreased from T0 to T1 in the tPBM group (t(18) = 2.30, p = .034; Supplementary Figure 1). No significant pre–post changes were observed for the other computational measures.

## 4. Discussion

Heterogeneity in response to non-invasive neuromodulation remains a central challenge, underscoring the need for markers that can characterize individual differences in clinical benefit. By applying computational modeling, this study provides preliminary evidence that task-derived computational markers may offer a comparatively scalable approach to characterizing individual differences in craving following non-invasive neuromodulation in subclinical drinkers receiving taVNS, tPBM, or their combination.

### Intervention and clinical-demographic factors

Among the intervention-related predictors, tPBM was associated with a greater likelihood of treatment response, consistent with the group-level craving reduction previously reported for tPBM in the source trial (Kwon et al., 2026), and prior evidence in individuals with opioid use disorder (Flora & Watson Huffer, 2024; Helali et al., 2025; Schiffer et al., 2021). In contrast, taVNS did not emerge as a stable predictor, consistent with limited evidence for its effects on alcohol craving (Kwon et al., 2026; Wang et al., 2021). Although taVNS can influence brainstem-cortical networks implicated in affective and stress regulation (C.-H. Liu et al., 2020; Badran et al., 2018), its role in alcohol craving outcomes warrants further investigation.

Baseline clinical-demographic characteristics also contributed to treatment response classification. Higher baseline craving severity was associated with a greater likelihood of response, consistent with prior rTMS findings in cocaine use disorder (Ghazi et al., 2025). This association may reflect greater opportunity for measurable improvement among individuals with higher initial craving.

Age and sex also appeared among the contributing variables. Sex/gender has been proposed as a relevant biological variable in brain-stimulation outcomes (Chmiel et al., 2026; Hanlon & McCalley, 2022), whereas age-related effects remain less clearly established (Leuchter et al., 2024; Slan et al., 2024). Therefore, the age and sex effects observed here should be interpreted as exploratory and require replication in larger, hypothesis-driven addiction-related studies.

### Computational markers

Beyond these clinical-demographic predictors, two computational modeling parameters also emerged as candidate predictors of treatment response, with higher loss aversion and lower risk sensitivity each associated with a greater likelihood of responder classification. These parameters characterize distinct aspects of value-based decision-making, reflecting the relative weighting of losses versus gains and valuation under known risk, respectively.

Prior evidence suggests that loss aversion may be relevant to alcohol– and substance-related outcomes: lower loss aversion has been associated with sustained heavy episodic drinking across emerging adulthood (Petzold et al., 2023) and with increases in substance use disorder symptoms over a one-year follow-up (Kräplin et al., 2020). One possibility is that individuals with greater loss aversion assign greater weight to the potential negative consequences of drinking relative to its anticipated rewards. If this tendency generalizes beyond monetary decision-making, it could support greater motivation for behavioral change and responsiveness to intervention.

Consistent with our a priori hypothesis, lower risk sensitivity—reflecting a less risk-seeking valuation profile—was associated with a greater likelihood of being classified as a treatment responder. Translated to alcohol-related decisions, this may indicate that the anticipated rewarding value of drinking is less likely to dominate when that reward is variable or not guaranteed. Prior alcohol-focused studies have shown that greater risk-taking propensity— the inverse of this profile—predicts subsequent relapse in AUD after clinical treatment (Yuan et al., 2024; Sasaki et al., 2023). Although derived from different risk-based paradigms, these findings converge in suggesting that model-derived risk valuation carries clinically meaningful information for alcohol-related outcomes.

Delay discounting did not show robust associations with treatment response, consistent with mixed evidence for its prognostic value in substance-use treatment (Exum et al., 2023).

For ambiguity sensitivity, our null finding is broadly consistent with prior work showing that day-to-day fluctuation, rather than baseline level, better predicts substance-use outcomes such as opioid and smoking relapse (Konova et al., 2020; J.-H. Lee et al., 2026); further replication is needed to determine whether this extends to neuromodulation-related craving outcomes.

Exploratory pre–post analyses showed a decrease in loss aversion in the tPBM group, consistent with a similar decrease in model-derived loss sensitivity reported by Todesco et al. (2022) during substance use treatment. However, as this analysis was exploratory and uncorrected for multiple comparisons, the finding is hypothesis-generating rather than evidence that changes in loss aversion underlie its baseline association with treatment response.

### Limitations

First, given the modest sample size and predictive performance, these computational parameters should be interpreted as preliminary indicators requiring validation in larger, independent cohorts. Second, the open-label design without a sham-control condition means that non-specific expectancy or placebo effects cannot be disentangled from stimulation-specific outcomes; future controlled trials are needed to establish whether baseline loss aversion and risk sensitivity specifically predict neuromodulatory efficacy. Third, our sample consisted of subclinical drinkers, which limits generalizability to clinical populations with severe AUD.

### Declaration of generative AI and AI-assisted technologies in the manuscript preparation process

During the preparation of this work, the authors used ChatGPT (OpenAI) for language editing and improving the readability of the manuscript. After using this tool, the authors reviewed and edited the content as needed and take full responsibility for the content of the publication.

### Data Availability Statement

The data that support the findings of this study are available from the corresponding author upon reasonable request, subject to institutional and ethical restrictions.

## Acknowledgements

This work was supported by the National Research Foundation of Korea (NRF) (RS-2024-00435727 and RS-2025-00516410 [to WYA], RS-2022-KH125035 [to YCJ, WYA, and JSC]), the Institute of Information & Communications Technology Planning & Evaluation (IITP) grant funded by the Korean government (MSIT) (RS-2026-25507282) [to WYA], the Korea Mental Health R&D Project, funded by the Ministry of Health & Welfare, Republic of Korea (HI22C0404) [to JSC], the Seoul National University Artificial Intelligence Graduate School Program (RS-2021-II211343) [to WYA], K-NeuroMind NEXUS: Neurocognitive-Affective Next-Generation Cross-modal Foundation Platform (RS-2026-25518987) [to WYA], and the BK21 FOUR Program (5199990314123) [to WYA]. The funders had no role in study design, data collection, data analysis, interpretation of the data, or preparation of the manuscript.

## CRediT authorship contribution statement

**Jungyoon Kang**: Conceptualization, Methodology, Formal analysis, Visualization, Writing – original draft. **Jung-Seok Choi**: Resources, Supervision, Funding acquisition, Writing – review and editing. **Young-Chul Jung**: Resources, Supervision, Funding acquisition, Writing – review and editing. **Minjae Kim**: Methodology, Formal analysis, Writing – review and editing. **Minkyung Park**: Investigation, Data curation, Project administration. **Manjae Kwon**: Investigation, Project administration, Data curation. **Yunesang Lee**: Validation, Writing – review and editing. **Dongil Chung**: Conceptualization, Methodology, Supervision, Funding acquisition. **Woo-Young Ahn**: Conceptualization, Methodology, Supervision, Resources, Funding acquisition, Writing – review and editing.

