## Supplementary file for "Baseline Predictors of Craving Reduction Following Non-Invasive Neuromodulation: A Computational Modeling Approach"

**Supplementary Materials**

**Supplementary Methods**

**Neuromodulation Conditions**

All participants self-administered their assigned intervention remotely for five weeks. The intervention schedule consisted of five sessions per week for five weeks, for a total of 25 sessions, with each session lasting 15 minutes. Before starting the intervention, participants attended an initial laboratory visit during which they were trained in device use, safety procedures, and adverse event reporting. Participants were also instructed to abstain from alcohol for at least 24 hours prior to each stimulation session. To ensure protocol compliance, research staff monitored device usage and intervention adherence through weekly check-ins.

The taVNS intervention group used the Healaon PRO device (Neurive Inc., Seoul, Republic of Korea). Following international consensus reporting standards for taVNS studies (Farmer et al., 2021), stimulation was delivered bilaterally to the cymba concha and cavum concha, using a bidirectional square-wave, non-burst waveform at 30 Hz and a pulse width of 200 μs. Intensity was individually titrated to a comfortable perceptible level, typically between 0.5 and 2.5 mA, and each session lasted 15 minutes.

The tPBM intervention group received transcranial near-infrared light using the iSyncMe device (iMediSync Inc., Seoul, Republic of Korea). Each 15-minute session delivered 850 nm light across 19 scalp channels (international 10–20 system) in three sequential phases: parietal/occipital stimulation at 10 Hz, whole-scalp stimulation at 15 Hz, and frontal stimulation at 18 Hz, each lasting 5 minutes. Posterior, whole-scalp, and frontal stimulation were selected to address key alcohol-related abnormalities: reduced alpha activity (Lew et al., 2020), widespread cortical dysregulation including elevated beta activity (Rangaswamy et al., 2002), and impaired prefrontal control over reward-driven behavior (Goldstein & Volkow, 2011). Additional details are provided in Kwon et al. (2026).

The combined taVNS + tPBM group received stimulation from both devices simultaneously within a 15-minute session, using the same parameters as the respective single-intervention groups.

**Reliable Change Index Calculation (RCI)**

The RCI was calculated following Jacobson and Truax (1991) as follows:

$$RCI = \frac{{PACS}_{T0}- {PACS}_{T1}}{\sqrt{2{({SD}_{T0}\sqrt{1-r_{xx}})}^{2}}}$$

where ${SD}_{T0}$ denotes the baseline standard deviation of PACS scores and $r_{xx}$ the reliability coefficient. ${SD}_{T0}$ was obtained from the present sample, and the PACS internal consistency estimate (Cronbach’s alpha = 0.91) from Kim et al. (2008) was used for $r_{xx}$.

**Supplementary Results**

**Exploratory pre-post changes in computational parameters**

Changes in computational parameters from T0 to T1 were examined in exploratory analyses within each intervention group. Loss aversion significantly decreased from T0 to T1 in the tPBM group (t(18) = 2.30, p = .034; Supplementary Figure 1), whereas no significant pre–post changes were observed for the other computational parameters in any intervention group. Because these analyses were exploratory and not corrected for multiple comparisons, the observed decrease in loss aversion should be considered preliminary.

**Supplementary Figures**

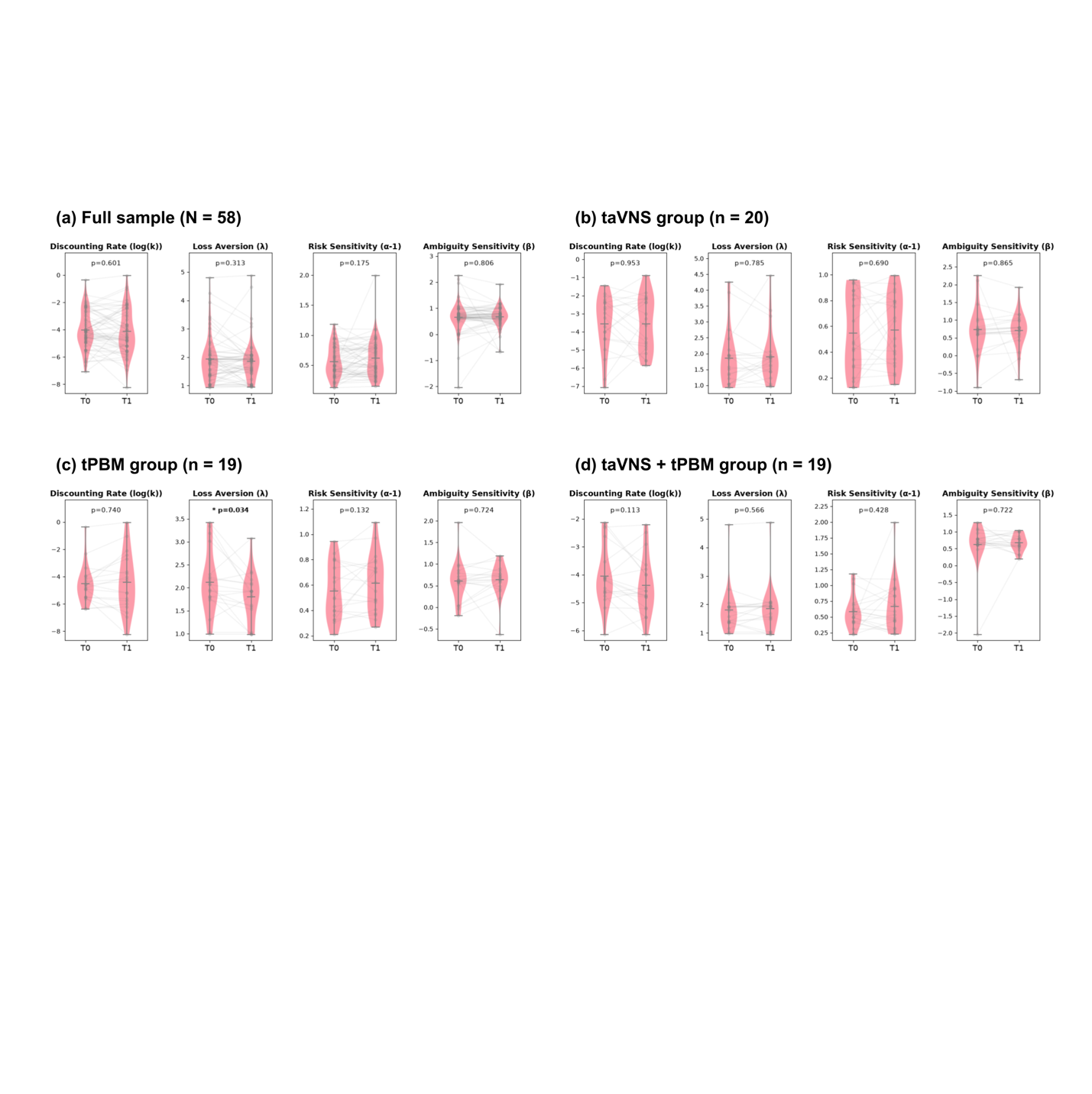

**Supplementary Figure 1**. Changes in computational parameters from T0 to T1 in the full sample and by intervention group. Pre–post changes in delay discounting, loss aversion, risk sensitivity, and ambiguity sensitivity are shown for the full sample and each intervention group. Lines connect paired observations from individual participants. Within-group changes were assessed using paired-samples t-tests. Exact p-values are shown for all comparisons, with statistically significant results highlighted in bold.
